# PhaseICA: Resolving Individual-Specific Macroscale Brain Propagations

**DOI:** 10.1101/2025.09.22.677524

**Authors:** Liang Ma, Qiang Li, Tulay Adali, Armin Iraji, Zening Fu, Vince D. Calhoun

**Affiliations:** Tri-Institutional Georgia State University/Georgia Institute of Technology/Emory University Center for Translational Research in Neuroimaging and Data Science (TReNDS), Atlanta, Georgia, USA; Department of Computer Science and Electrical Engineering, University of Maryland, Baltimore County, Baltimore, Maryland, USA; Department of Computer Science, Georgia State University, Atlanta, Georgia, USA

## Abstract

Understanding brain function requires knowing not only which regions fluctuate together, but also how their activity propagates across space and time. Yet current fMRI representations may not provide concise, accurate, subject-specific descriptions for propagation patterns, limiting insights into brain organization and dysfunction. Here, we introduce PhaseICA, a framework combining higher-order complex source separation with individualized wave reconstruction. Simulations with known ground truth demonstrate more accurate recovery of complex mixtures of overlapping standing and traveling waves than conventional approaches. Applied to human resting-state fMRI, PhaseICA resolves a reproducible repertoire of macroscale waves and reconstructs their subject-specific patterns. These representations improve cross-session identification, enhance crystallized cognition prediction and schizophrenia classification, while revealing behavior- and symptom-associated changes in propagation amplitude and timing. By resolving individual propagation dynamics within functional activities, PhaseICA extends brain mapping beyond co-fluctuation representations toward a comprehensive description of how nonlinear coordination unfolds across space and time.

## Introduction

Brain function is organized through coordinated functional activities across distributed systems. Using functional magnetic resonance imaging (fMRI), researchers have mapped blood oxygenation level-dependent (BOLD) co-fluctuations into several large-scale functional networks ^1,2^, with their topology and temporal coupling highly associated with cognition, behavior and psychiatric status^3-10^. Traditionally, we describe brain activity as fixed spatial maps modulated by separate time courses ^7,11-15^. This space-time separation favors the characterization of synchronous or standing-like fluctuations across brain regions, but cannot parsimoniously represent the spatial traveling flow. Different from standing activity, spatial traveling flow involved a nonlinear relationship between space and time, with coordinated regions engaged at systematically different phases. The route and timing of a flow can carry information that static maps do not retain ^16-18^. A whole-brain account of phase-ordered dynamics could therefore extend conventional representations of functional activity from static co-fluctuation maps to descriptions of how nonlinear coordination unfolds across space and time.

Traveling waves provide a concise representation for travelling activities. Electrophysiological and optical recordings have revealed planar, spiral and recurrent traveling waves in visual, motor and association cortices, with varying direction and geometry ^19-22^. These micro- and mesoscale observations show the biological existence of traveling propagation, but their restricted observed scopes cannot establish its whole-brain structure. fMRI provides complementary coverage and could also reveal macroscale, slow propagations or variance-dominant spatiotemporal modes^18,23-25^. These findings show the presence of macroscale propagation but do not determine whether the recovered modes have been separated accurately and independently. Currently, complex principal component analysis (CPCA) ^18,26^ has therefore been used to extract dominant traveling modes from Hilbert-transformed analytic signals, which encode spatial amplitude and relative phase. However, the accuracy of these recovered latent waves remains unclear. CPCA relies on second-order covariance to optimize wave patterns. It may consequently combine overlapping standing and traveling patterns, displace lower-variance waves, or be influenced by periodic and aperiodic BOLD activities^27^. Since empirical fMRI lacks the ground truth, reproducing findings or explained substantial variance does not establish the recovered accuracy. Moreover, current studies have generally characterized only a small number of dominant modes. When multiple latent flows are mixed within a single dominant mode, the effects from specific propagation patterns could be diluted or obscured by contributions from other constituent flows. Resolving these questions requires more accurate separation beyond covariance, and a principled determination on component granularity, as well as simulated validation against known ground truth.

Besides recovered accuracy, establishing underlying associations of these propagations also require individual-level precision^28,29^. Traditionally, individual differences in conventional network topology support subject identification ^30^ and behavioral association^3,4^. Similarly, previous studies have also linked the regional lag structure to cognition, arousal, development and behavior^23,31-33^, and identified altered timing in schizophrenia^34^. Building an accurate wave-phenotype association not only require the recovered accuracy on common space but also requires subject-specific amplitude and phase-delay maps expressed within this common space. If so, individual wave representations should improve the performance of downstream tasks, e.g. cross-session individual identification, behavioral prediction and psychiatric status, because more features are encoded in travelling waves than pure standing waves. Accurately localizing phenotype-associated amplitude and phase-delay variances within macroscale wave patterns could help identify regional phenotype contributions and help investigate underlying neural coordination.

Here we introduce PhaseICA, a pipeline that integrates population-level wave separation with subject-specific reconstruction. PhaseICA contains a full procedure of individual wave separation based on higher-order statistics than a second-order covariance. Specifically, it converts fMRI time series into analytic signals and applies the established complex entropy bound minimization (CEBM) algorithm^35^ to estimate population independent sources. CEBM captures higher-order distributional information through entropy bounds constrained by expectations of nonlinear measuring functions. Permutation-based parallel analyses and stability analyses across repeated trails determinate the component granularity. Finally, a spatially-constrained extension of CEBM enables a prior-based wave individualization to obtain subject-specific amplitude and phase-delay maps.

We tested two linked hypotheses. First, we assumed a higher-order complex decomposition should recover the complicated mixture of standing and traveling sources more accurately than standing-wave decompositions, and a second-order statistic wave decomposition. Second, an individual representation, from a higher-order statistic decomposition pipeline, should improve downstream task performance than pervious alternatives. To verify the first hypothesis, we evaluated recovery accuracy against known ground truth in 2D simulations, involving different combinations of overlapping waves (space: standing/traveling, time: periodic/aperiodic). PhaseICA showed the highest accuracy in the recovery of complicated simulated sources. Then we made task validations on resting-state fMRI from 995 Human Connectome Project participants (HCP)^36^ and 311 Functional Imaging Biomedical Informatics Research Network (FBIRN) participants^37^, including 151 with schizophrenia. PhaseICA improved simulated source recovery and identified reproducible macroscale waves. Their individualized representations improved cross-session identification, selectively enhanced cognitive prediction and increased schizophrenia classification accuracy.

These representations also provided a biological perspective on multi-region asynchronous interactions across different individuals. Crystallized cognition was associated with wave amplitude in left inferior frontal language cortex and relative phase in adjacent regions, suggesting that acquired verbal abilities are reflected in both regional response magnitude and the timing of language-related regions within distributed activity patterns. In schizophrenia, PhaseICA revealed altered phase relationships involving the thalamus, cerebellum and caudate within a wave spanning prefrontal-thalamic-cerebellar circuitry, associated with negative-symptom severity. This organization is consistent with the disrupted circuit coordination proposed by the cognitive dysmetria hypothesis^38^. By distinguishing regional amplitude from relative timing within individual waves, PhaseICA offers a framework for investigating how distributed coordination relates to cognitive ability and psychiatric dysfunction.

## Results

To facilitate understanding, we provide a 1D visualization illustrating the concept of standing and traveling waves (Fig. 1a). Standing waves *Y*(*x, t*) = *A*(*x*)*B*(*t*) *cos*[*φ*(*x*)] *cos*[*ω*(*t*)] can be represented by separable spatial and temporal terms, whereas traveling waves *Y*(*x, t*) = *A*(*x*)*B*(*t*) *cos*[*φ*(*x*) + *ω*(*t*)] encode location-dependent phase and therefore couple space and time. Standing waves are a special case of travelling waves when *sin*[*φ*(*x*)]*sin*[*ω*(*t*)] = 0.

**Fig. 1.**
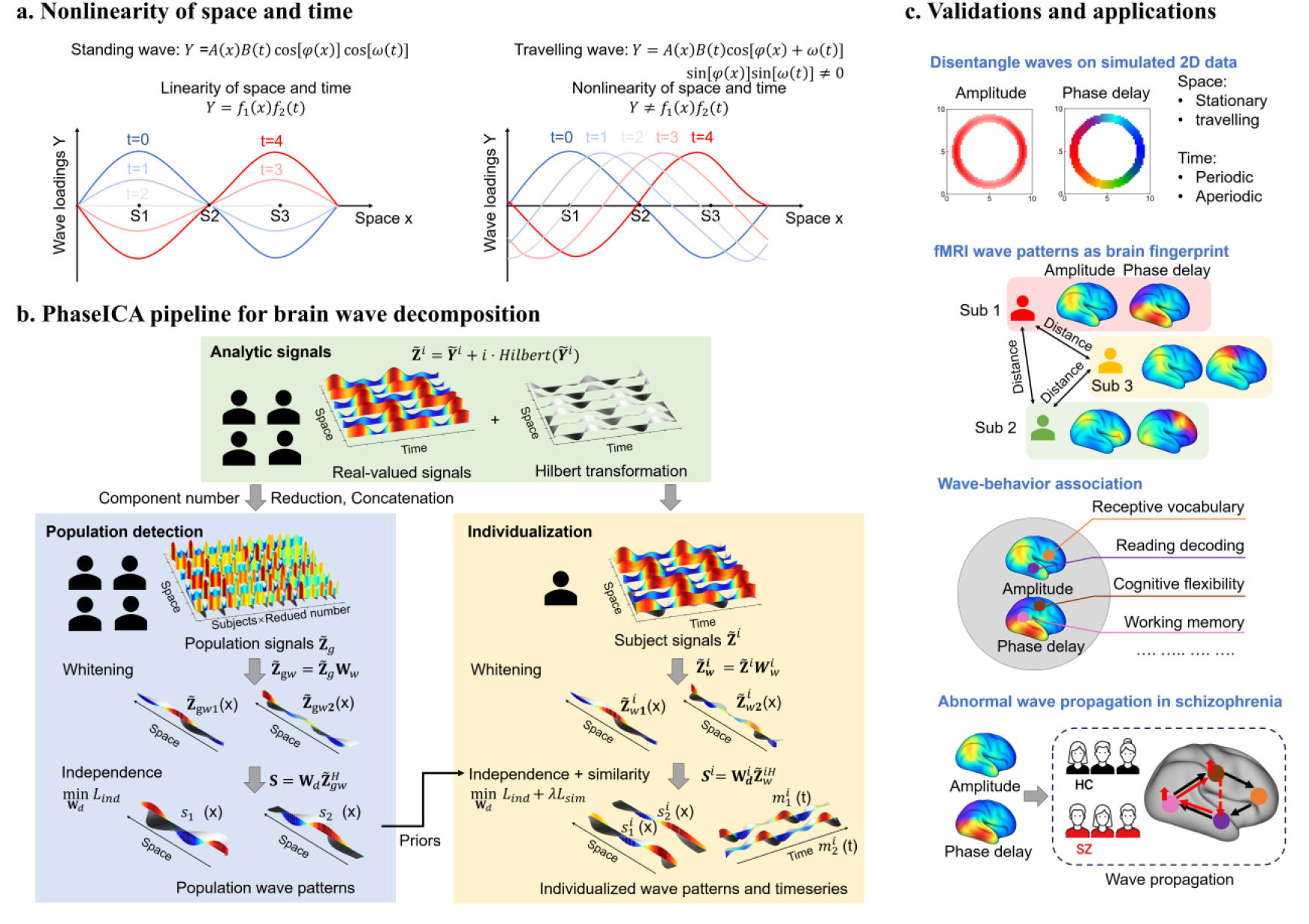
PhaseICA workflow, validation and applications. **a:** Separable space-time coupling in standingwaves and nonlinearity in traveling waves. **b:** PhaseICA workflow. For wave detection, each real-valued signal is first converted to an analytic signal by Hilbert transformation, followed by dimensionality reduction. The reduced number is determined by parallel analysis and stability analysis. Reduced subject signals are concatenated, whitened and separated by CEBM. For individualization, each analytic signal is whitened and decomposed by spatially constra ined CEBM, which jointly optimizes component independence and similarity to the population priors. **c:** Validation of wave recovery on simulated 2D data and downstream analyses of brain fingerprinting, wave-behavior associations and schizophrenia classification.

PhaseICA aims to address the individual traveling representation problem within macroscale BOLD activities through higher-order source separation in the complex domain (Fig. 1c). We came up with two linked questions: whether PhaseICA could accurately recover wave representations under complicated wave conditions and whether wave representations are robust within individual and improve the performance of downstream tasks. We first assessed source recovery, noise robustness, model-order selection and individualization against known ground truth in simulations.

We then examined reproducibility and granularity in human fMRI, followed by cross-session subject identification, behavioral prediction and schizophrenia classification. Further analyses were conducted to identify brain regions in the wave propagation that substrate behavioral or psychiatric outcomes. In the Supplementary Materials ‘Mathematical proof’, we provide a theoretical analysis to support accurate separation of traveling waves using higher-order source separation.

### Higher-order complex statistics improve recovery of complicated wave mixtures

Since empirical fMRI does not provide source-level ground truth, the accuracy of traveling wave recovery has to be evaluated by simulation with known ground truth. We generated four wave classes on a 50 × 50 grid crossed standing or traveling structure with periodic or aperiodic temporal dynamics. A fifth, mixed condition contained all four classes and provided the most stringent test of overlapping sources. Based on the simulation, we are able to test whether the higher-order-statistic optimization adopted by PhaseICA could improve the recovery accuracy than CPCA based on the second-order statistic (covariance).

In a representative mixed simulation (Fig. 2a, b), three traveling waves (W1-W3) propagated clockwise around separate rings. Forward propagation direction follows the negative spatial-phase gradient. Phase velocities, defined as −[∂ω(t)/∂t]/[∂φ(x)/∂x], were 3.34, 1.26 and 1.31 m s^−1^ for W1, W2 and W3, respectively. W4 and W5 waves were standing waves fluctuated within fixed regions. W2, W3 and W5 had periodic temporal dynamics. The angular frequencies (∂ω(t)/∂t) of W2 and W3 were 0.063 and 0.052 rad s^−1^, respectively. W1 and W4 were aperiodic, with frequencies that varied around their base values. Based on this ground truth, we compared CEBM results with those from two real-valued methods, principal component analysis (PCA) and spatial independent component analysis (sICA), as well as the complex-valued method, CPCA (Fig. 2c, d). CEBM separated all five sources while preserving each wave amplitude and phase-delay distributions. Instead, comparison methods are likely to mix multiple wave patterns within one component.

**Fig. 2.**
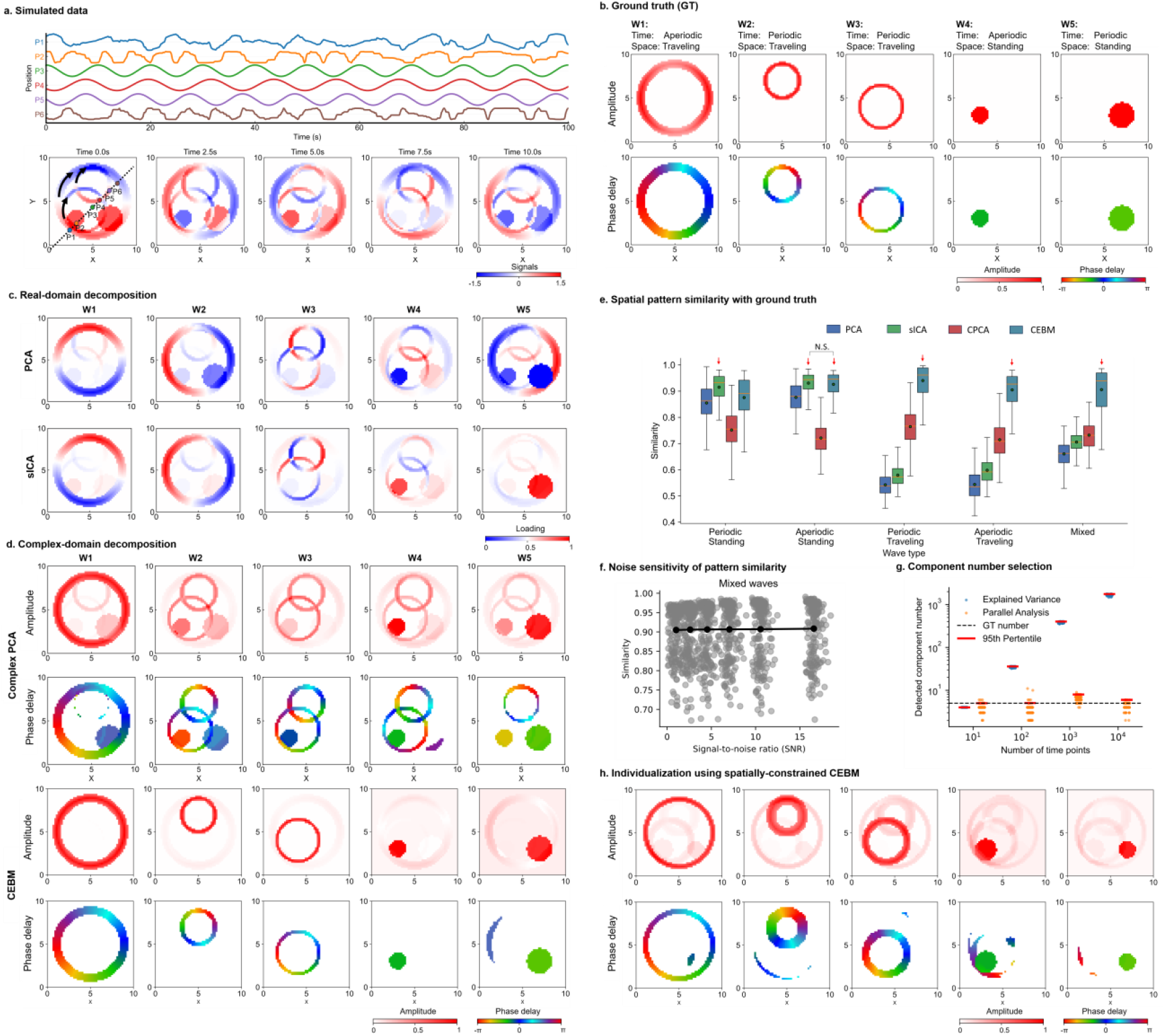
Recovery of simulated standing and traveling wave patterns. **a:** Temporal signals and representative spatial frames from a mixed simulation containing five waves. **b:** Ground-truth amplitude and phase-delay maps for three traveling and two standing waves. **c:** Components recovered by the real-valued methods PCA and sICA. **d:** Amplitude and phase-delay maps recovered by CPCA and CEBM. For c and d, amplitude maps were normalized to the corresponding ground-truth maximum, and phase was displayed where normalized amplitude exceeded 20% of the maximum. **e:** Similarity between recovered components and ground-truth waves across the four wave classes and mixed condition in 200 simulated datasets. The red arrow indicates the best performance in each situation. N.S. indicates FDR-corrected p>0.05 in a two-sample t-test. **f:** Similarity between CEBM components and ground-truth waves across SNRs from 0.1 to 15. Gray points denote individual simulations, and the black line denotes the mean. **g:** Model order selected by 95% explained variance or parallel analysis as the number of time points increased. The dashed line denotes the ground-truth model order of five, and red markers denote the 95th percentile. **h:** Example individualized components after varying ring bandwidths and radii while holding wave centers fixed . The example shows high pattern similarity (0.94) with individual GT. GT, ground truth; SNR, signal-to-noise ratio.

We evaluated the performance of different methods across 200 simulations. There are randomized wave centers, radius, bandwidths, phase velocities and angular frequencies, and additional Gaussian noise (σ=0.3). For each method, recovery was quantified as the mean maximum similarity between ground-truth waves and estimated components. Among the tested methods, sICA performed best for standing periodic waves (0.91 ± 0.05; CEBM, 0.87 ± 0.08). Both sICA and CEBM achieved the highest performance for standing aperiodic waves without significance (0.94 ± 0.04; CEBM, 0.93 ± 0.04, two-sample t test, p=0.56). CEBM achieved the highest similarity for traveling periodic waves (0.94 ± 0.06), traveling aperiodic (0.90 ± 0.07) and mixed waves (0.91 ± 0.06). In these last three complicated conditions, CEBM significantly outperformed PCA, sICA and CPCA (two sample t test, FDR-corrected P < 0.05; Fig. 2e). It shows that higher-order statistical, complex-valued separation could gain more advantageous when wave sources are more complicated.

To investigate the sensitive of high-order statistic decomposition to the data quality, we varied the standard deviation of the Gaussian noise from σ = 0.1 to 0.6, producing signal-to-noise ratios (SNRs) from 0.1 to 15 in the mixed-wave simulated condition. The CEBM decomposition maintained a mean ground-truth similarity of approximately 0.90 across SNRs, indicating limited sensitivity to unstructured noise (Fig. 2f). CEBM computation time and memory usage scale approximately linearly with the temporal and spatial dimensions of the data (see Supplementary Fig. S1).

Accurate separation also depends on model order. Previous testing is based on known ground truth order. Next, we develop a procedure to decide the model order that most closely approximates the ground truth order. The model order is first determined by the parallel analysis (PA) against the permutations and finally filtered by ICASSO stable analysis (seen Method “Wave number determination”). We compared a conventional 95% explained -variance criterion with a 95% PA number. For each participant, the PA number was defined as the total number of empirical eigenvalues exceeding their rank-matched thresholds from 200 structured permutations. The 95th percentile of these individual PA numbers could achieve a population model order more closed with ground truth (see Methods, ‘Wave number selection’). With the ground-truth order fixed at five, the model order selected by explained variance increased sharply as the number of time points rose from 10 to 10,000. Parallel analysis instead selected 5, 5, 6 and 5 components at 10, 100, 1,000 and 10,000 time points, respectively (Fig. 2g). Finally stable components are all 5 after stable filtering (ICASSO Iq>0.9) across repeated trails.

Population-level recovery does not ensure that participant-level reconstruction preserves individual variation. We therefore generated 200 individual datasets from the mixed -wave simulation with Gaussian noise (σ = 0.3), varying ring bandwidths and circle radii by -50% to +50% while holding wave centers fixed. Spatially constrained CEBM maintained correspondence with the population priors while retaining the simulated variation (Fig. 2h). The individualized components achieved a similarity of 0.91 ± 0.05 to their individual ground truth.

### A highly reproducible macroscale wave repertoire

Having evaluated source recovery in simulations, we next examined the spatial organization of macroscale waves in empirical fMRI. We applied PhaseICA to resting-state fMRI from 100 unrelated participants in the HCP young adult dataset, divided equally into discovery and validation cohorts (n=50 each). Population-level decomposition identified 12 waves, with model order set to the 95th percentile of subject-level PA numbers. All components showed greater reproducibility between cohorts than CPCA (PhaseICA mean=0.92, CPCA mean=0.84, **Supplementary Fig. S2**).

We summarize their phase-delay maps and amplitude profiles across the seven Yeo functional networks^1^, subcortical regions and cerebellum (Fig. 3a). Wave amplitude and phase delay were normalized (see Methods “Population-level wave detection”). The fine-grained wave repertoire showed network-specific spatial propagation. Specifically, each wave showed dominant amplitude in one to several functional networks or subregions, with lower-amplitude regions generally exhibiting relative phase offsets. The traveling index (TI) of waves ranged from 0.11 for W10 to 0.65 for W3, indicating a continuum from modest to pronounced traveling characteristics.

**Fig. 3.**
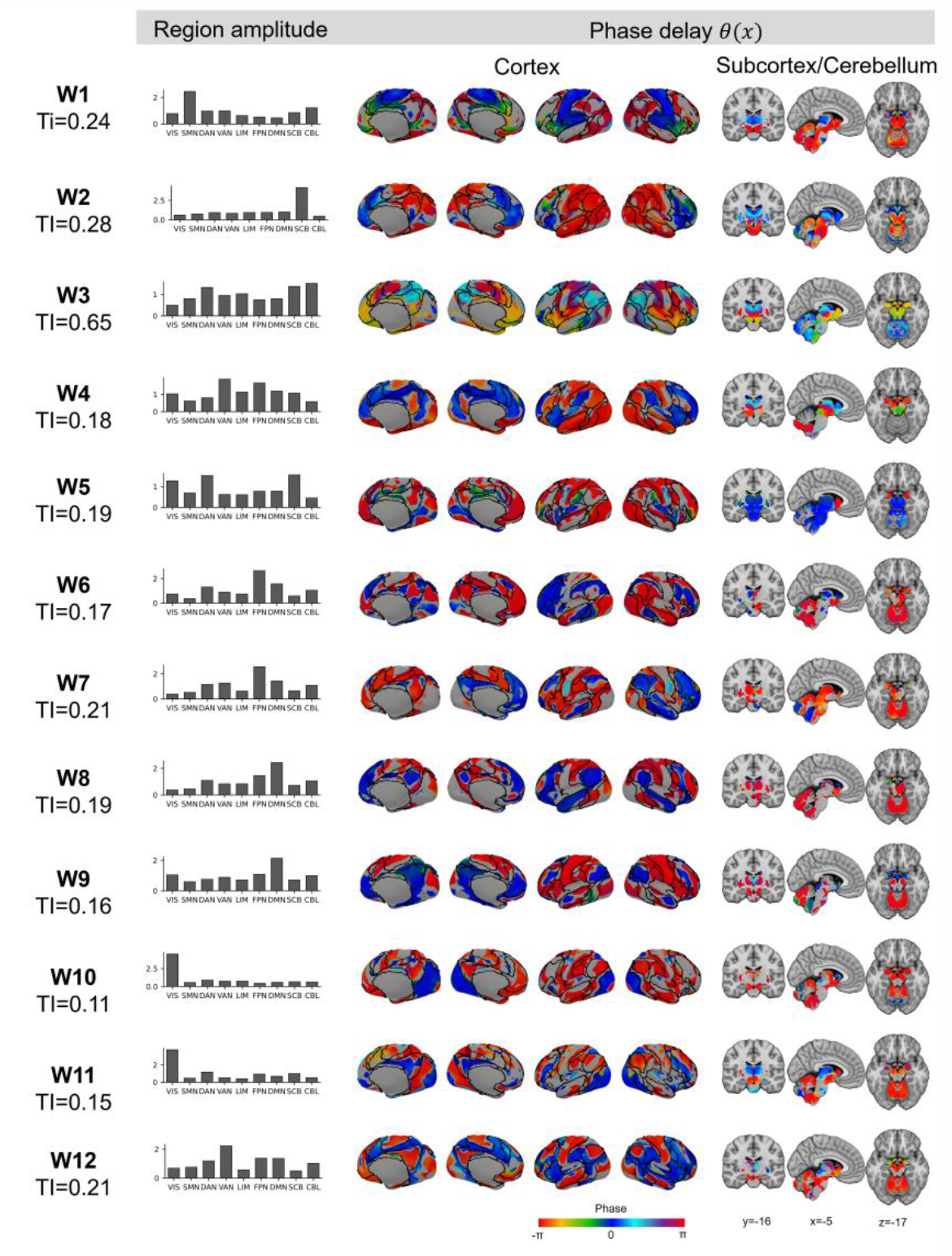
12 fine-grained macroscale waves resolved from PhaseICA pipeline. Phase-delay maps and mean amplitudes across nine brain systems for 12 fine-grained waves. Amplitudes were averaged within the seven Yeo cortical networks 1, subcortex and cerebellum. Each wave’s whole-brain mean amplitude was normalized to 1, and global phase was rotated so the maximum-amplitude voxel had phase 0. Low-amplitude region (<0.5) is masked. TI, traveling index. High TI values indicate strong traveling expression. DAN, dorsal attention network. DMN, default mode network; FPN, frontoparietal network; LIM, limbic; SMN, somatomotor network; VAN, ventral attention network; VSN, visual network; SBC, subcortex; CBL, cerebellum.

### Individual propagation patterns provide robust brain fingerprints

Population-level reproducibility is insufficient for precision analyses. A robust representation of macroscale brain waves should capture individual-specific propagation patterns that remain stable within participants while differing across participants. We first inspected and quantified spatial variation within these macroscale waves. As an example of individual variation, the population pattern of W3, a strongly traveling wave, begins with high amplitude in the cerebellum near phase 0 (**Fig. 4a, b**). Its phase evolution then traced an ordered route through bilateral ventral striatal regions (π/2), ventromedial prefrontal cortex (π/2∼π), insular cortex (π/2∼π), and postcentral gyrus (3/2π), precuneus (3/2π) and medial superior frontal gyrus (3/2π) (**Fig. 4a**). Individual wave maps show significant deviations from this population pattern in both regional amplitude and phase delay. As indicated by light yellow circles (**Fig. 4b**), participant 133019 showed an increased amplitude in bilateral occipital lobes and altered phase delay in postcentral gyrus. Participant 499566 showed an increased amplitude in the right medial primary motor cortex and bilateral occipital lobes, accompanied by altered phase delay in bilateral ventral striatal regions. Participant 756055 also showed lower amplitude and altered phase delay in the bilateral ventromedial prefrontal cortex. Across all 12 PhaseICA waves, the mean similarity was 0.65 ± 0.04 across different subjects and 0.80 ± 0.03 between two sessions from the same subject (mean ± s.d.; **Fig. 4c**).

**Fig. 4.**
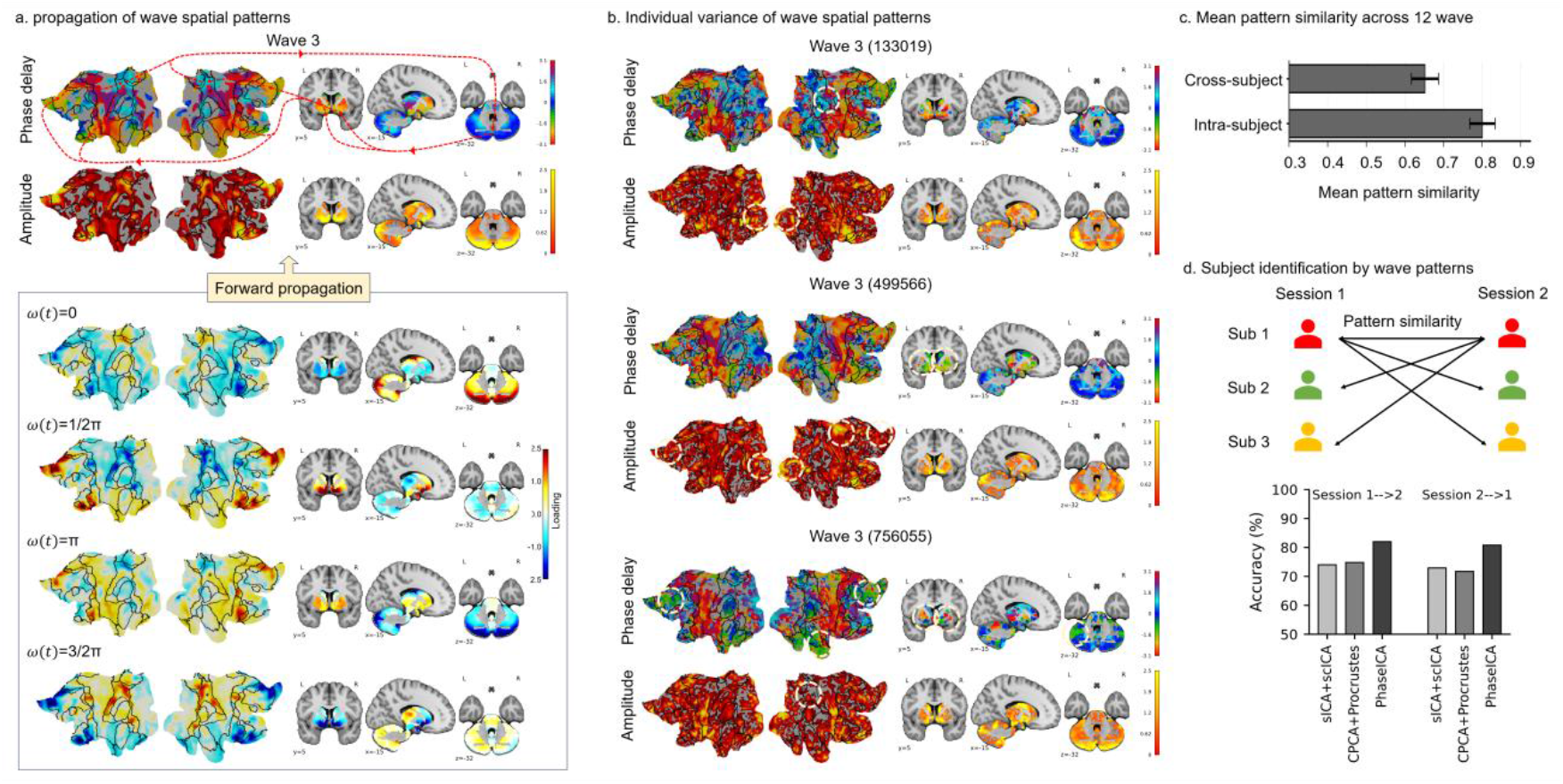
Individual wave patterns as a robust brain fingerprint. **a:** Population W3 phase-delay and amplitude maps (top) and its phase progression with spatial loadings at phases ω(t) = 0, π/2, π and 3π/2 (bottom). Red lines and arrows indicate forward propagation through high-amplitude subcortical and cortical regions. b: Individual W3 phase-delay and amplitude maps for subject 133019, 499566 and 756055. Dashed circles mark individual deviations from the population pattern. c: Mean cross-subject and within-subject pattern similarity across 12 waves. d: Cross-session identification accuracy across three spatial representations (sICA + scICA, CPCA+Procrustes and PhaseICA).

We next tested whether this combination of within-subject stability and cross-subject variability improved cross-session subject identification. We compared subject-specific PhaseICA maps with two commonly used pipelines (see Supplementary Materials for comparison methods “Comparison pipelines”). The network-topography pipeline uses population sICA followed by spatially constrained ICA (sICA + scICA); the second-order wave pipeline uses CPCA followed by Procrustes alignment (CPCA+ Procrustes). All methods used 12 subject-specific spatial maps per session and were evaluated in 991 participants with eligible scans in both sessions. For each cross-session participant pair, we averaged the similarities between the 12 corresponding maps with equal weight (Methods, ‘Subject identification using wave patterns’). Each participant was matched to the candidate in the opposite session with the highest mean similarity. PhaseICA achieved the highest accuracy (82% and 80.8%) for both session 1-to-2 and session 2-to-1 matching (**Fig. 4d**). These exceeded the corresponding accuracies of sICA + scICA (74.0% and 72.9%) and CPCA (74.8% and 71.7%). Under matched model order, PhaseICA individual wave patterns provided the most idiosyncratic brain fingerprints among the three representations.

### Higher-order complex representations selectively enhance wave-behavior associations

Subject identification establishes individual specificity but not phenotypic relevance. We therefore asked whether the higher-order complex representations recovered by PhaseICA improved behavioral prediction beyond fixed network topography (sICA + scICA) and second-order complex representations (CPCA + Procrustes). We compared sICA + scICA, CPCA + Procrustes and PhaseICA pipelines using the same number of components. The three approaches represented fixed network topography, second-order wave representations and higher-order wave representations, respectively. Ridge regression predicted HCP crystallized and fluid cognition composites as well as their 7 cognitive subtest scores using ten repeats of tenfold cross-validation (Methods, ‘Ridge regression for cognitive prediction and schizophrenia classification’; Fig. 5a).

**Fig. 5.**
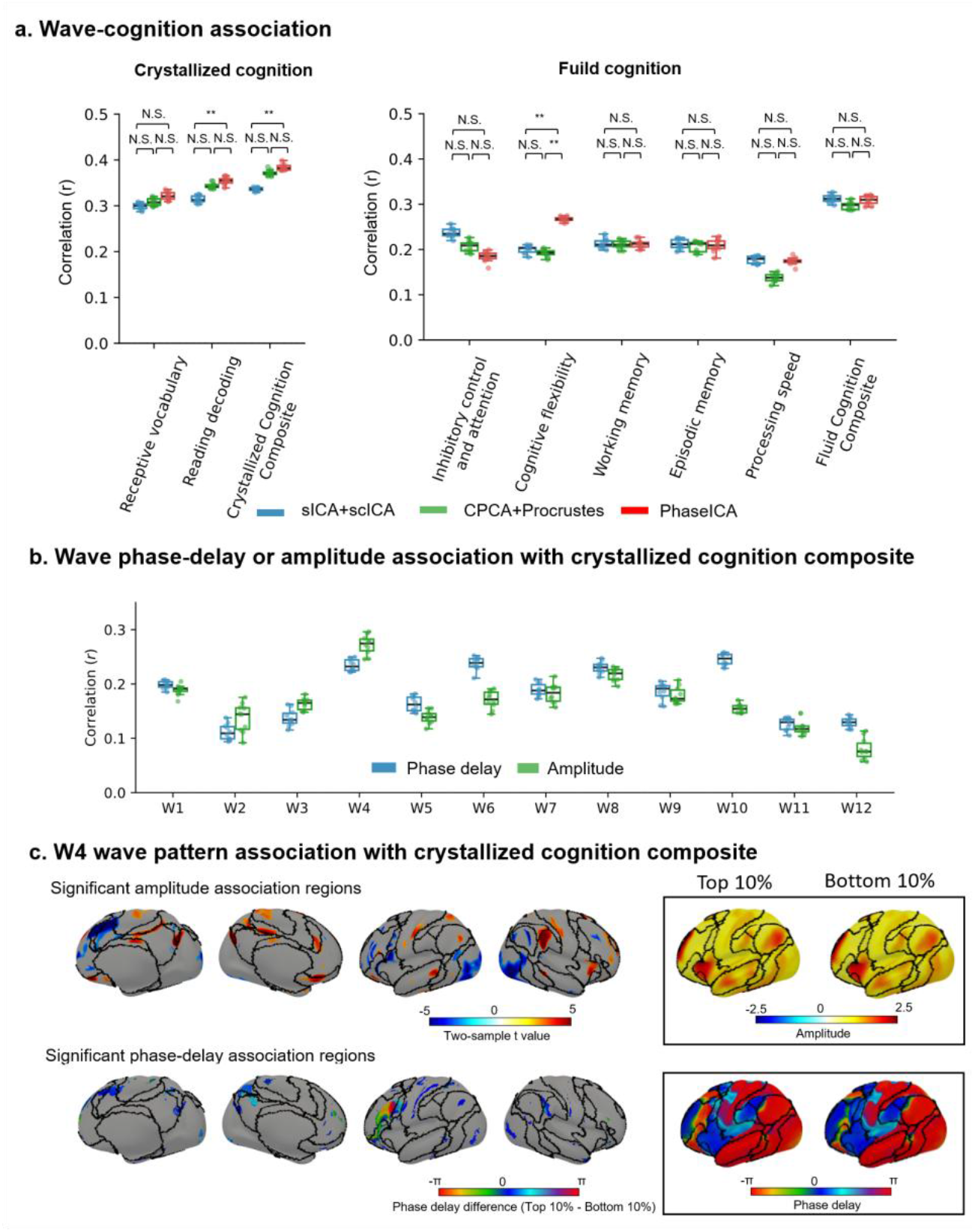
Higher-order complex representations selectively enhance brain-behavior associations. **a:** Observed-predicted correlations (r) for crystallized and fluid cognition composites and their 7 cognitive subtest scores from ridge regression using sICA + scICA, CPCA + Procrustes or PhaseICA representations (n = 979 participants). The significance of performance differences between methods was tested under 10000 subject-level permutations with p-values corrected across behaviors. **b:** Wave-specific prediction of crystallized cognition using each PhaseICA wave’s phase-delay or amplitude maps. Boxplots in a and b summarize ten repeats of tenfold cross-validation. **c:** Spatial associations between W4 amplitude or phase delay and crystallized cognition. Voxel-wise general linear models were adjusted for age, sex and head motion. Row 1 shows significant amplitude-associated regions (two-sample t test, t(974), FDR-corrected P < 0.05). Row 2 shows significant regions with phase-delay-associated regions (joint F-test, f(2, 973), FDR-corrected P < 0.05). Color maps show mean phase-delay differences between participants in the highest 10% and lowest 10% behavioral deciles. The left inset shows mean amplitude and phase-delay maps for these two groups. N.S., not significant. ^**^, significant.

The advantage of PhaseICA was selective across different cognitive domains. PhaseICA produced a higher observed-predicted correlation (r=0.384 ±0.006) for the crystallized cognition composite than network topography (r=0.336 ± 0.004, one-sided paired permutation test, 10000 permutations, FDR-corrected p <0.05) as well as second-order wave representations (r=0.371 ± 0.005, one-sided paired permutation test, 10000 permutations, p=0.269). Predictions on “receptive vocabulary” and “reading decoding” scores also show a similar performance, with a higher correlations from PhaseICA ( r=0.321 ± 0.008 and 0.354 ± 0.008 ) than those obtained using the sICA+scCIA methods (0.299± 0.006 and 0.308 ± 0.007, one-sided paired permutation test, 10000 permutations, FDR-corrected p=0.150 and p<0.05) as well as the CPCA + Procrustes method (0.314± 0.007 and 0.344 ± 0.006, one-sided paired permutation test, 10000 permutations, FDR-corrected p=0.269 and p=0.269). Higher-order representation could encode more individual variance related with crystallized cognition than the other two methods. However, PhaseICA showed no prediction advantage for fluid cognition composite. Except for “cognition flexibility”, wave associations with fluid cognition composite and its four subtest score analyses showed no gains from PhaseICA representations. It indicates specific cognition-related variances has already fully encoded in standing or second-order representations.

To identify which higher-order-statistic wave representations were most associated with crystallized cognition, we estimated the performance from separate phase-delay and amplitude models. As shown in **Fig. 5b**, W4 provided the strongest amplitude-based prediction (mean r ± s.d., 0.28 ± 0.02) and one of the strongest phase-delay predictions (0.23 ± 0.01). W4 linked high-amplitude expression in bilateral frontoparietal and ventral attention networks near 0 phase-delay, lower-amplitude expressions in default mode and limbic networks, visual cortex and subcortex (**Fig. 5c**). Voxel-wise general linear models adjusted for age, sex and head motion localized these associations. Higher crystallized cognition was associated with lower W4 amplitude near phase 0 in the left Broca area (two-sample t test, t(974), FDR-corrected P < 0.05). A broader set of regions adjacent to the left Broca area showed significant phase-delay associations (joint F-test, f(2, 973), FDR-corrected P < 0.05), indicating shifts in phase delay near the left Broca area in participants with higher crystallized cognition. These findings mapped the regional amplitude and phase-delay features associated with crystallized cognition within a macroscale wave derived from the PhaseICA pipeline.

### Phase-resolved wave patterns improve schizophrenia classification and localize altered propagation

We next tested whether the representational advantage of PhaseICA generalized to psychiatric status. We used resting-state fMRI from 321 FBIRN participants, including 164 participants with schizophrenia (SZ) and 157 healthy controls (HCs). Our model order procedure filtered 10 stable waves for the FBIRN dataset. FBIRN wave patterns were matched their HCP counterparts with a mean pattern similarity of 0.64. This correspondence supported partial cross-dataset reproducibility despite differences in cohort and acquisition sampling rates (HCP: TR=0.72s, FBIRN: TR=2s). A slightly higher model order may be detected by a higher sample-rate data, because more detailed differences could be present across components. We then compared the SZ classification performance across three representations (sICA + scICA, CPCA + Procrustes and PhaseICA) with the same model order, and the same ridge regression model. Fig. 6a shows PhaseICA achieved the highest classification accuracy (mean ± s.d., 0.778 ± 0.005), exceeding sICA + scICA (mean ± s.d., 0.766 ± 0.009, one-sided paired permutation test, 10,000 permutations, p=0.24) and significantly outperforming CPCA + Procrustes (0.735 ± 0.006, one-sided paired permutation test, 10,000 permutations, p<0.05). Thus, adding higher-order phase-resolved information improved case-control separation beyond second-order complex patterns.

**Fig. 6.**
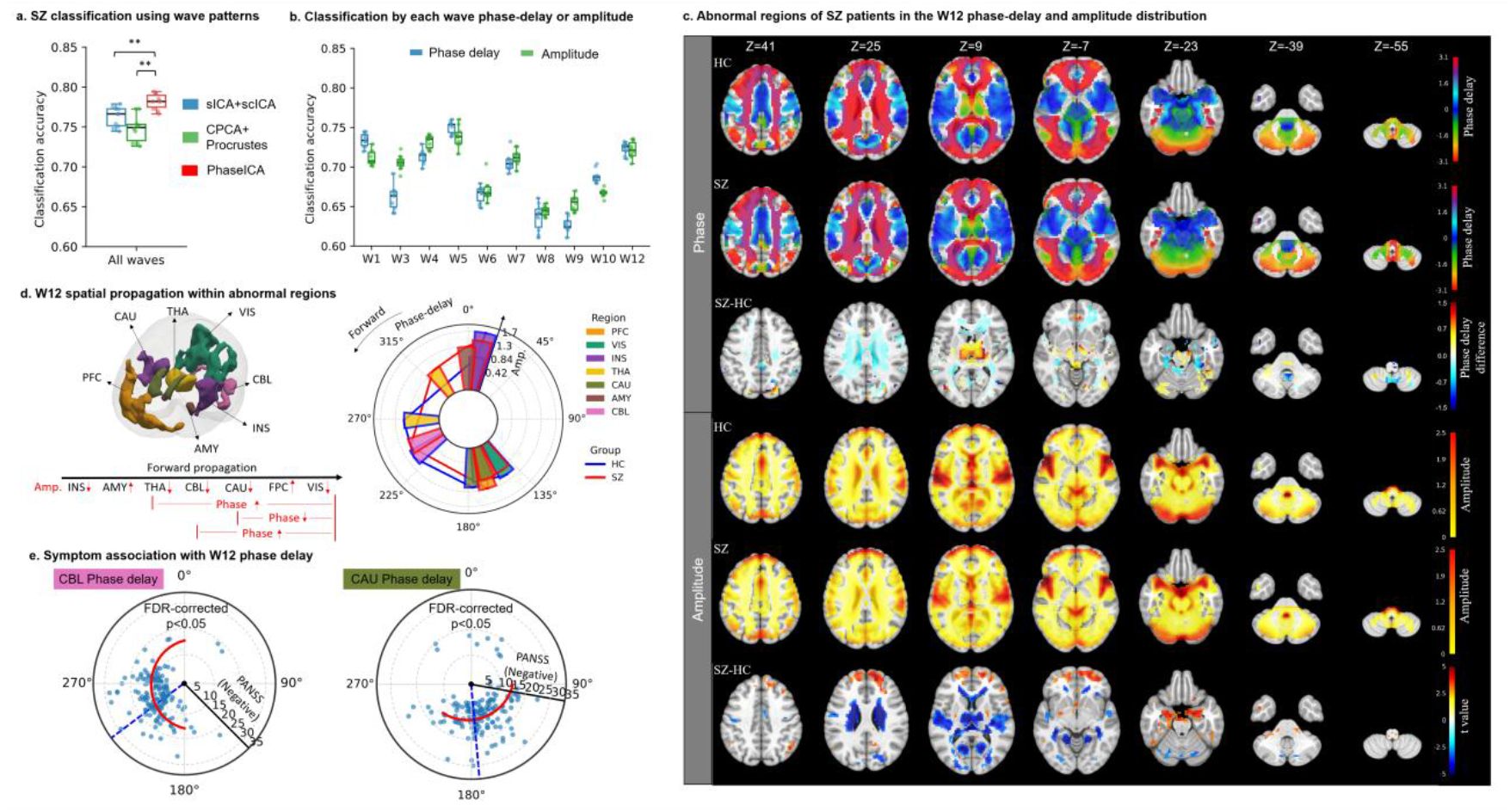
PhaseICA wave patterns improve schizophrenia classification and localize altered pathways. a: SZ classification accuracy using individualized spatial patterns from sICA + scICA, CPCA + Procrustes or PhaseICA (151 SZ, 160 HCs). **b:** SZ classification accuracy using the phase-delay or amplitude map of each PhaseICA wave. The same cross-validation splits were used across all three representations to ensure paired comparisons. **c:** mean W12 phase-delay maps within groups are shown in rows 1-2, followed by the SZ-HC phase-delay difference in row 3. Phase-delay difference regions are masked by the significance of GLM (joint F-test, f(2, 298), FDR-corrected P < 0.05). Rows 4-5 show mean amplitude maps within groups, and row 6 shows significant SZ-related voxels via general linear model (two-sample t test, t(299), FDR-corrected P < 0.05). **d:** 3D localizations of significant W12 regions and its propagation profiles within two groups. Arrows indicate SZ-HC differences in amplitude and phase-delay maps. The polar map shows regional mean amplitudes and phase delays. **e:** Associations between cerebellar/caudate phase delays with PANSS negative-symptom scores among SZ patients. Red curves show fitted associations, blue dots represent each SZ patient and blue dashed lines denote mean HC phase delays. PFC, prefrontal cortex; VIS, visual cortex; INS, insula; THA, thalamus; CAU, caudate nucleus; AMY, amygdala; CBL, cerebellum . Amp., Amplitude.

After observing this classification advantage, we asked which waves, features and regions built this association. We tested the performance from separate phase-delay and amplitude models, with W1, W4, W5 and W12 features exceeding 0.70 mean accuracy (Fig. 6b). W12 was chosen because of its propagation from the insula through the amygdala, thalamus, cerebellum and caudate to prefrontal and visual cortices (Fig. 6c, d). High-amplitude regions (prefrontal, thalamus, and cerebellum) in W12 overlap with the prefrontal-thalamic-cerebellar circuitry in previous SZ findings^38^. We filtered brain regions in which there are significance associations between W12 amplitudes or phase delays and SZ status using voxel-wise general linear models (FDR correction

P < 0.05; **Fig. 6c**). We found, relative to HCs, SZ patients showed shifted phase delays in the thalamus, cerebellum and caudate along forward propagation. They showed lower W1 amplitude in the insula, thalamus, cerebellum, caudate and visual cortex, together with higher amplitude in prefrontal cortex and amygdala. After removing small isolated areas, we manually delineated 7 abnormal regions to characterize SZ differences in W12. We further confirmed significant differences (two-sample t test, t(299), FDR-corrected p<0.05) in mean regional phase delay and amplitude between SZ patients and HCs (**Fig. 6d**). Within the SZ group, cerebellar and caudate phase delays were associated with PANSS negative-symptom scores (joint F-test, f(2, 298), FDR-corrected P < 0.05; **Fig. 6e**). Participants whose phase delays were closer to the HC reference had lower negative-symptom severity. The PhaseICA representation that improved classification also localized schizophrenia-related variation to specific timing and amplitude changes along a specific functional pathway.

## Discussion

In this study, we presented the PhaseICA pipeline to characterize phase organization beyond that captured by traditional ICA-based pipelines. We validated two linked assumptions in macroscale standing and traveling wave analysis. First, we validated against known ground truth in simulated 2D data. Higher-order statistical decomposition in our pipeline recovered complicated waves more accurately than traditional standing-wave decomposition or second-order complex decomposition. Second, PhaseICA resolved fine-grained macroscale waves spanning standing-like to strongly traveling patterns. The downstream tasks derived from these fine-grained waves have shown improved cross-session identification, selectively improved behavioral associations and increased schizophrenia classification accuracy over matched alternative methods. Regional analyses identified timing or amplitude features within specific waves that were strongly associated with behavioral measures and psychosis findings. Together, these results support our pipeline based on higher-order statistics as a method for more accurately characterizing macroscale nonlinear dynamics.

PhaseICA provides a parsimonious, individualized description of macroscale standing and traveling activities at the whole-brain scale with more accuracy. Macroscale traveling wave here refers to the propagation of traveling BOLD hemodynamic activities. Under the temporal positive/negative gradients of phase time series, there is a forward/backward propagation of hemodynamic response across multiple regions following the evolving sequence of phase delays. Conventional functional networks emphasize zero-lag correlation among multiple regions, whereas a PhaseICA wave jointly describes its high-amplitude nodes, their relative delays and the sequence linking them. This representation separates two contributions often conflated in associations with phenotype: response magnitude and propagation phase/timing. Our pipeline also developed procedures to determine the effective number of waves, which was validated by simulation data. It resolved a fine-grained repertoire of macroscale neural flows rather than only a few variance-leading modes ^18,33^. This separation allows phenotypic associations to be attributed to specific flows, although their biological origin requires further validation.

PhaseICA recovers these macroscale waves without prescribing the periodic or aperiodic fluctuation^27^, spatial autocorrelation ^39^, wave geometry ^22,23^, and a linear state-transition model ^40-42^. It does not require a spatial reduction from functional networks or sliding-window length to characterize the nonlinear interaction among multiple regions^43-45^. The pipeline also accommodates purely standing-like patterns. We wish to reduce the dependence on assumptions that can strongly shape functional dynamics. The propagation direction, geometry, frequency and temporal evolution could be characterized after source separation. The pipeline also retains several assumptions. We assumed that population-level, complex-valued sources are statistically independent, and individual sources largely overlap with population sources but with individual variations. Also, signal preprocessing will influence the recovered repertoire, but it is testable and remains adaptable to specific scientific questions.

Crystallized cognition analysis illustrates how accurate wave separation can improve behavioral associations. Previous work has shown that vocabulary and reading recognition performance is associated with functional connectivity within a coordinated, distributed language-semantic system, involving left inferior frontal and temporal language regions and broader frontoparietal and default-mode networks^46-48^. PhaseICA W4 wave pattern overlaps with these regions. W4 linked high-amplitude expression in the inferior frontal gyrus and anterior and medial prefrontal cortex around phase-0, with delayed, amplitude-reduced responses in anterior temporal lobes and lateral parietal cortex, and middle frontal gyrus. Beyond characterizing the behavioral association with co-fluctuation activities, our individual wave pattern provides the propagation timing of the left middle frontal gyrus within this system play a significant factor. Accurate travelling representation could extend brain-behavior analysis from functional connectivity analysis to the investigation on how their amplitude and timing are organized within a coordinated system.

The analysis of schizophrenia patients showed a similar advantage for higher-order statistical representation. Previously, dysfunction of prefrontal-thalamic-cerebellar circuitry has been implicated in cognitive dysmetria and schizophrenia^38^. Earlier work identified abnormal activation or connectivity within this system^49,50^, but those measures could not determine whether altered coupling reflected changes in response strength or relative response order. PhaseICA wave representation showed altered W12 propagation within this circuitry of SZ patients, involving amplitude changes in the insula, amygdala, thalamus, cerebellum, caudate, prefrontal cortex and visual cortex, and shifted response order in the subcortex, e.g. thalamus, cerebellum and caudate. Within the schizophrenia group, cerebellar and caudate phase delays closer to the healthy-control reference were associated with lower PANSS negative-symptom severity, consistent with a recent negative symptom association in a broader psychosis population ^51^. Although current findings do not establish a detailed biological interpretation, a more precise characterization of propagation patterns may help clarify mechanisms underlying schizophrenia. Integrating these patterns with regional maps of cell types, neurotransmitter systems, or receptor distributions could help reveal underlying mechanisms for competing hypotheses.

By resolving overlapping macroscale waves and reconstructing subject-specific amplitude and phase-delay maps, PhaseICA provides a general framework for characterizing whole-brain propagation in fMRI. Although developed for BOLD signals, its mathematical formulation is not restricted to this modality. Applications to EEG, MEG and local field potentials could resolve traveling patterns at higher temporal resolutions and across different spatial scales, helping relate macroscale fMRI propagation to electrophysiological dynamics in neuronal populations. Cross-modal analyses may further clarify how waves emerge, propagate and interact across scales, although modality-specific preprocessing, source localization and validation will be required. The framework may be adapted to multivariate spatiotemporal data containing overlapping propagating sources. Potential applications include metabolic dynamics, climate systems and transportation networks. Although these extensions remain to be tested, they suggest that higher-order complex decomposition may offer a general approach for separating propagation patterns in complex systems.

## Methods and datasets

### Datasets

#### 1. Simulated wave dataset

We generated four wave conditions that crossed spatial organization (standing versus traveling) with temporal dynamics (periodic versus aperiodic), together with a mixed condition; implementation details are provided in Code Availability. Each sample contained five waves on a 50 × 50 grid with 0.2-m spacing. Traveling waves (phase velocity ≠ 0) propagated along annular trajectories whose radius, location and width varied, whereas standing waves (phase velocity = 0) remained within a fixed circular region. Traveling-wave phase velocities ranged from 0.038 to 1.256 m s^−1^. For periodic waves, frequency *F* ranged from 0.006 to 0.2 Hz. For aperiodic waves, instantaneous frequency comprised a constant baseline in the same range plus a smooth stochastic deviation constrained to remain positive:Δ*F*(*t* + 1) = α Δ*F*(*t*) + *ε*(t). The deviation followed a first-order autoregressive process equivalent to a discrete-time Ornstein–Uhlenbeck process^53^. We used *α* = *e*^−1/0.15^, with the Gaussian innovation *ε* scaled by 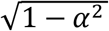 . Each realization contained 1,000 time points sampled at 1-s intervals. Spatial wave amplitude was determined by annulus radius and width, 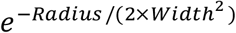, whereas the temporal amplitude B(t) was fixed at 1. Spatial phase θ(x) was determined by the wave region and phase velocity, and temporal phase ω(t) was obtained from instantaneous frequency. The resulting wave signal was*Y* = A(x)B(t)cos [θ(x) + ω(t)], and the five wave signals were summed in a space-time array to form a sample.

Wave locations and spatial overlap were randomized to model propagation across partially overlapping functional organizations. Annulus location and width, circle radius, phase velocity and angular frequency varied across samples. Gaussian noise with spatially varying scale was added to assess noise sensitivity. The signal-to-noise ratio was calculated as 10*log*_10_(∑s*ig*n*al*^2^ /∑ *noise*^2^).

To evaluate individualization, we generated 200 variants of each simulated population reference. Wave location, phase velocity and angular frequency were held constant, whereas annulus width and circle radius were independently increased or decreased by 10–50% relative to the reference.

#### 2. Human Connectome Project dataset

We used the publicly available Human Connectome Project Young Adult (HCP-YA) S1200 dataset, which comprises approximately 1,200 participants aged 22–35 years^36^. Data were acquired on a customized 3 T Siemens Connectome Skyra scanner with high-performance gradients. The protocol included high-resolution structural MRI, resting-state fMRI and behavioral assessments. Resting-state fMRI used a multiband gradient-echo echo-planar imaging sequence. Each participant completed two sessions with two runs per session. Each run comprised approximately 1,200 time points (repetition time, 720 ms; echo time, 33.1 ms; flip angle, 52°; 2.0-mm isotropic resolution; 72 slices; multiband factor, 8). The two runs in each session used left-to-right and right-to-left phase encoding.

HCP fMRI data first underwent minimal preprocessing^54^, including gradient-distortion correction, head-motion correction, intensity normalization, EPI distortion correction, co-registration and generation of grayordinate time series in CIFTI space. Multimodal surface matching (MSMAll) was used to support accurate cross-participant surface registration^55^. Further postprocessing was performed using XCP-D ^56^, with explicitly specified parameters. This included nuisance regression using 24 motion regressors (the six rigid-body motion parameters, their temporal derivatives, and the squared terms of both sets), followed by temporal bandpass filtering at 0.01-0.15 Hz and spatial smoothing with a Gaussian kernel of 6 mm full width at half maximum. Runs with mean framewise displacement (FD) >0.3 mm were excluded ^57^. To avoid removing global fluctuations potentially relevant to the propagation patterns of interest, global-signal regression was not applied. 995 participants (532 females) remained after quality control. Subject identification included 991 HCP participants with eligible scans in both sessions after motion-based exclusion. Four of the 995 participants were excluded because eligible repeated scans across the two sessions were unavailable. Population wave priors were estimated from 100 participants who were also included in the identification cohort. Population-level wave estimation used 100 independent participants (22-35 years; 54 females), divided equally between discovery and validation sets (N=50 each). Runs were whitened separately before population decomposition, and the principal findings were evaluated in both sets.

We examined wave associations with crystallized and fluid cognition composites with 7 their cognitive subtest scores using NIH Toolbox assessments ^58,59^. Specifically, the crystallized Cognition Composite combined receptive vocabulary and reading decoding scores, reflecting acquired verbal knowledge and skills.receptive vocabulary was measured by word-picture matching and reading decoding through oral word recognition. The Fluid Cognition Composite summarized performance across inhibitory control and attention, cognitive flexibility, working memory, episodic memory and processing speed. Inhibitory control and attention were assessed using the Flanker task, which requires responding to a target while ignoring distracting stimuli. Cognitive flexibility was measured by switching sorting rules in the Dimensional Change Card Sort task. Working memory was assessed through recalling and reordering items in the List Sorting task. Episodic memory was measured by reproducing picture sequences, whereas processing speed was assessed through rapid visual pattern comparisons. Subject with any missing labels are excluded (n=16) in the cognition analysis.

#### 3. Functional Imaging Biomedical Informatics Research Network dataset

The Functional Imaging Biomedical Informatics Research Network (FBIRN) Phase III dataset comprises 362 adults: 176 individuals with schizophrenia or schizoaffective disorder and 186 healthy controls^37,60^. Participants were recruited at seven US academic medical centers and scanned on seven 3 T MRI systems. Diagnoses were established using structured clinical interviews based on the SCID for DSM-IV-TR, and patients were clinically stable for at least two months. Symptom severity was assessed with the Positive and Negative Syndrome Scale^37^. Acquisition followed the standardized FBIRN traveling-phantom protocol to reduce inter-site variation^61^. Resting-state fMRI used a harmonized gradient-echo echo-planar imaging sequence. Each scan comprised 162 volumes acquired over 5 min 24 s (repetition time, 2,000 ms; echo time, 30 ms; flip angle, 77°; field of view, 220 × 220 mm; voxel size, 3.4 × 3.4 × 4 mm). Participants kept their eyes closed, remained awake and minimized movement.

The fMRI data were processed using the NeuroMark pipeline^62^. The first 16 volumes were discarded, followed by head-motion correction, EPI distortion correction, slice-timing correction and nonlinear registration to Montreal Neurological Institute (MNI) space. Quality control required fMRI scans to have at least 120 remaining volumes, with the head motion <3° rotation and <3 mm translation at any time point (n=51). Images then underwent head-motion regression, band-pass filtered at 0.01-0.15 Hz and smoothed with a 6-mm full-width at half-maximum Gaussian kernel. No frames were censored. After filtering, 311 participants remained, with 151 schizophrenia patients.

### PhaseICA pipeline

PhaseICA first estimates population-level complex spatial sources and then derives spatially constrained subject-specific counterparts. Throughout this section,bold uppercase symbols denote matrices, bold lowercase symbols denote vectors and unbolded symbols denote scalars. The superscript H denotes the Hermitian transpose, ∥·∥_2_ denotes the Euclidean norm and Re(·) denotes the real part. K denotes the number of wave components.)

### Population-level wave detection

For each participant, let **Y** ∈ ℝ^*V*×*T*^ denote the real-valued data matrix, where V is the number of spatial locations and T is the number of time points. Each matrix was z-standardized across all spatial locations and time points.

#### Analytic representation

The Hilbert transform was applied to the time series at each location to retain instantaneous amplitude and phase^63^:

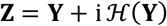

Here, i is the imaginary unit and *H*(·) is the Hilbert transform. The first and last 10 time points were excluded to reduce finite-length boundary artefacts.

#### Complex reduction and whitening

For each participant, dimensionality reduction and whitening used the eigenvectors and eigenvalues of temporal covariance matrix 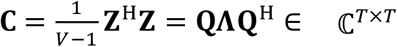. The K leading eigenvectors and eigenvalues ( 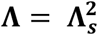, where **Λ**_***s***_contains singular values from the SVD decomposition) were retained by fast singular value decomposition^64^. A whitening matrix transformed its covariance matrix into an identity matrix to obtain the whitened spatial matrix: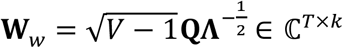

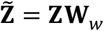

Thus, 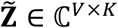 has identity covariance ( 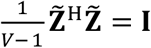, **I** is identity matrix) across spatial locations. Parallel analysis determined K, as described below. Participant -level whitened matrices were concatenated along their reduced dimensions. A second K-dimensional complex reduction and whitening step yielded the population matrix 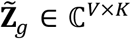.

#### Complex independent component analysis

Complex entropy bound minimization (CEBM) ^35^ was applied to the transposed population matrix to estimate population-level spatial sources:

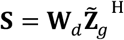

Here, **S** ∈ ℂ^*K*×*V*^ contains K complex spatial sources, and **W**_*d*_ ∈ ℂ^*K*×*K*^ is the demixing matrix. CEBM minimizes

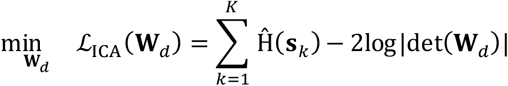

For source **s**_*k*_, Ĥ(**s**_*k*_) is the upper bound of the maximum entropy of **s**_*k*_. Each candidate entropy bound was constrained by expectations of nonlinear measuring functions, including the weighted-linear-combination and elliptical families. For a complex source **s**_*k*_, *x* = Re[**s**_*k*_] cos(*θ*) + Im[**s**_*k*_] sin(*θ*), the weighted-linear-combination family used the statistics 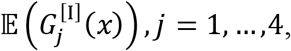 where 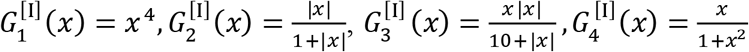. The elliptical family used the radial statistics 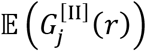 *j* = 1,2, where *r* = |**s**_*k*_| and 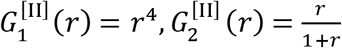. The fourth-order functions capture kurtosis and radial tail weight, whereas the even and odd rational functions capture broader tail behavior and distributional asymmetry. The tightest candidate entropy bound was retained for each source, enabling CEBM to accommodate noncircular, sub-Gaussian, super-Gaussian, symmetric and skewed distributions. Optimization used the projected conjugate-gradient procedure described in the original CEBM optimization^35^.

The population wave pattern was obtained using the ICASSO procedure^65^ by running CEBM 20 times. After ICASSO, each qualified (ICASSO Iq<0.9) wave’s mean amplitude was normalized to 1, and its maximum-amplitude voxel was assigned phase 0.

### Subject-specific wave estimation

Each participant underwent the same analytic-signal construction, K-dimensional reduction and complex whitening. The subject-specific sources were estimated as

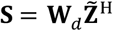

where **S** ∈ ℂ^*K*×*V*^. The rows of Wd are the component-specific demixing vectors. The population source matrix **R** ∈ ℂ^*K*×*V*^ provides one spatial reference **r**_*k*_ for each component. Individualization balances CEBM independence with correspondence to these references:

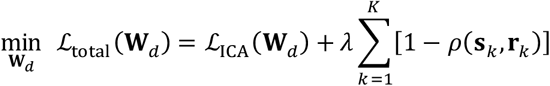

The spatial-prior weight was used to balance the loss from component independence and spatial correspondence with priors. Complex cosine similarity was defined as

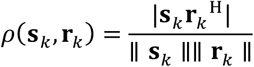

The demixing matrix was initialized by the least-squares projection of R onto the participant’s whitened data: 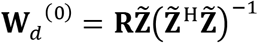 . The objective was optimized row by row using Wirtinger derivatives with respect to the conjugate demixing vector^66^. The independence and similarity gradients were normalized by the scale at the first iteration before combination:

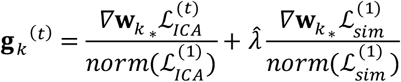

At each iteration, the combined gradient was projected onto the tangent space of the unit -norm constraint:

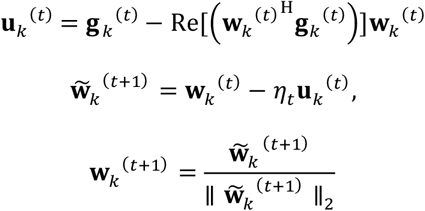

Optimization was performed separately for two statistical families: the weighted -linear-combination family and the elliptical family. For the optimization of each family, the learning rate *η*^*t*^ was initialized at 0.04 and halved after three consecutive iterations without a decrease in the objective. Optimization stopped when this criterion recurred after *η*^*t*^ fell below 0.005 or when the maximum iteration count (1000) was reached.

After convergence, the general demixing matrix **D** and its mixing matrix **M** were

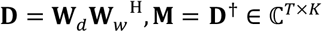

The k^th^ column of **M** provided the complex time course **m**_*k*_ corresponding to spatial source **s**_*k*_. The phase-delay and amplitude of individual sources were then aligned to their population references: 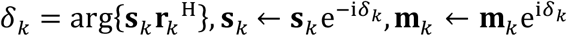. Finally, each source was divided by its mean spatial amplitude, with the reciprocal scaling applied to its time course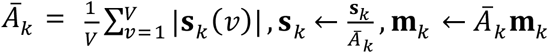. These coupled transformations preserved each component’s reconstructed real-valued contribution.

### Wave number determination

We used parallel analysis to determine the supported number of components and ICASSO to assess their stability.

#### Parallel analysis

Independent circular shifts were applied to the time series at each spatial location. This procedure preserved marginal distributions and temporal autocorrelation while disrupting cross-location covariance ^67,68^. Eigenvalues were recomputed for each null realization. At each rank, the 95th percentile of the null distribution defined the threshold. The subject-specific PA number was the total number of empirical eigenvalues exceeding their rank-matched thresholds. The 95th percentile of these subject-specific counts (the nearest integer) was used as the approximate population wave number.

#### ICASSO stability analysis

Complex-valued ICASSO stability analysis was applied to test component stability^65^. CEBM was repeated 20 times from random initializations for a fixed whitened population matrix. Components were grouped by hierarchical clustering of pairwise complex spatial similarities. Stability was quantified using the ICASSO Iq index, which ranges from 0 to 1. Higher values indicate more reproducible estimates. Stable population components (ICASSO Iq >0.9) were represented by central representatives of stable clusters.

### Wave interpretation of PhaseICA components

For component k, complex spatial source **s**_*k*_ and complex time course **m**_*k*_ define its real-valued contribution at spatial location *v* and time *t*:

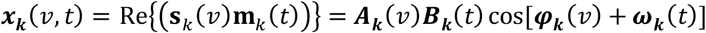

The spatial amplitude and phase terms were

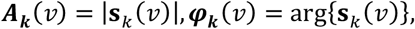

and the temporal amplitude and phase terms were

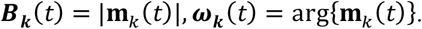

The real and imaginary parts of the spatial loadings represent the loading pattern at phase ***ω***_***k***_(*t*) = 0 and ***ω***_***k***_(*t*) = *π*/2 . The real-valued loading at an arbitrary phase ***ω***_***k***_(*t*) was 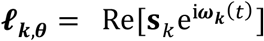. Each population component was rotated by 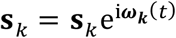 to place the maximum-amplitude voxel at phase zero.

### Quantification of spatial propagation patterns

#### Component complex similarity

Similarity between two complex spatial components **s**_*i*_ and **s**_*j*_was the magnitude of their normalized Hermitian inner product:

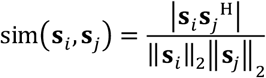

where ‖**s**_*i*_‖_2_ is the norm of **s**_*i*_ . This measure ranges from 0 to 1 and is invariant to global phase. Before comparison with a real-valued map, the complex component was rotated to maximize cosine similarity between the real map and the component’s real part.

#### Traveling index

The traveling index^18,69^ was the inverse condition number of the matrix formed by the real and imaginary spatial loadings **L**_*k*_ = [Re[**s**_*k*_], Im[**s**_*k*_ ]]:

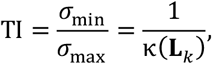

Values near 0 indicate nearly collinear real and imaginary patterns, as expected for a standing or standing-like component. Values near 1 indicate linearly independent quadrature patterns and a stronger traveling-wave character.

### Subject identification using wave patterns

Subject identification used spatial-pattern similarity across two repeated sessions. Each method provided the same number of subject-specific maps per session (12 in this study), with components matched across sessions. For every participant pair across sessions, we calculated the absolute complex similarity between corresponding maps. The 12 component similarities were then averaged to obtain an overall participant-pair similarity (matrix shape: [participant, participant]). Each participant in the query session was matched to the participant in the reference session with the highest mean similarity. A match was correct when both represented the same individual. Identification accuracy was the percentage of correctly matched participants. We performed matching in both directions and reported accuracy separately for each direction.

### Ridge regression for cognitive prediction and schizophrenia classification

Ridge regression tested how well subject-specific spatial patterns predicted cognitive scores or schizophrenia status^70^. Ridge regression model was described as below. Let **X** ∈ ℂ^*N*×*P*^ denote the complex feature matrix for N participants and P spatial features, and let **y** ∈ ℝ^*N*^ denote the real-valued target. Cognitive outcomes were continuous. Schizophrenia status was coded as 1 for schizophrenia and 0 for healthy control. The complex ridge estimator minimized

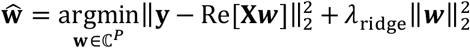

Both real and imaginary features contributed to the model, whereas predictions remained real-valued, *λ*_ridge_ is the penalty parameter. Because P exceeded N, estimation used the dual representation:

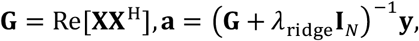

The dual weights are 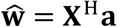, and prediction was obtained by 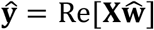.

Performance was evaluated using ten-fold cross-validation repeated ten times with the ridge penalty *λ*_ridge_ = 1. Subject-level out-of-fold predictions were concatenated in a cross-validation. For the HCP, the subjects were removed in the training fold split which has family relationship with out-of-fold subjects. Subject-specific wave patterns have been normalized before the prediction. Cognitive prediction was quantified by the Pearson correlation between observed and predicted scores. For schizophrenia classification, predicted scores were converted to diagnostic labels using a threshold of 0.5, and performance was evaluated using classification accuracy. Besides all-wave features jointly, we evaluated wave-specific phase-only or amplitude-only features separately. Phase-only features were represented by the unit phase **s**_*k*_/| **s**_*k*_|, whereas amplitude-only features were | **s**_*k*_|.

The same component number, spatial number and cross-validation procedure were used for the representations across different pipelines. The sICA + scICA pipeline implemented in GIFT toolbox, combined group spatial ICA^71^ with spatially constrained individualization^72^, as used in NeuroMark-style network delineation^62,73^. The CPCA + Procrustes pipeline^32,74^ matched individual CPCA components to their population counterparts by orthogonal Procrustes alignment. PhaseICA pipeline combined population CEBM with spatially constrained CEBM individualization. Implementation settings for the comparison pipeline are described in the **Supplementary Materials**.

Differences in predictive performance across methods were assessed using one-sided, subject-level paired permutation tests with 10,000 permutations. All methods were evaluated using 10 repetitions of 10-fold cross-validation with identical data partitions. Predictive performance was quantified as the mean Pearson correlation between predicted and observed behavioral scores for HCP behavioral prediction, and mean classification accuracy for schizophrenia classification, both averaged across the 10 repetitions. The observed test statistic was defined as the performance difference between methods. For each permutation, paired predictions (cognition) or participant-level accuracies (schizophrenia) were randomly exchanged between methods with a probability of 0.5 for each participant, preserving the same exchange assignment across repetitions. Performance differences were then recalculated to construct the empirical null distribution. One-sided p-values were estimated as the proportion of permuted differences greater than or equal to the observed difference, using a plus-one correction. Statistical significance was determined at p < 0.05, with Benjamini-Hochberg FDR correction applied to multiple comparisons across different behaviors.

### Spatial associations of wave amplitude and phase with target variables

Spatial associations were tested independently at each location using general linear models (GLM) ^75^. Let target ***y*** ∈ ℝ^*N*^ contains continuous cognitive scores for HCP or binary diagnostic status for SZ (HC = 0; SZ = 1). The covariate matrix **C**_cov_ ∈ ℝ^*N*×*Q*^ included sex, age and head motion for behavioral analyses, with scanner-site indicators additionally included for SZ analyzes. N denotes the number of participants; Q is the number of covariate columns. Participants with missing target or covariates were excluded.

#### 1) Amplitude associations

At location *v*, the amplitude model was

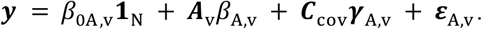

At a single location, ***A***_v_ ∈ ℝ^*N*×1^ contains one amplitude measurement per participant. The intercept *β*_0A,v_ and amplitude coefficient β_A,v_ are scalars. The nuisance coefficient vector ***γ***_A,v_ ∈ ℝ^*Q*×1^ multiplies ***C***_cov_ on the right. The null hypothesis *H*_0_: *β*_A,v_ = 0 and two-sided t statistic were 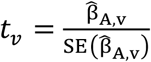. The residual degrees of freedom were N-Q-2, (N, Q for sample number and covariate number). Positive and negative coefficients indicate associations between greater amplitude and higher or lower target values, respectively.

For HCP, the amplitude coefficient represents the expected change in cognitive score per one-unit increase in wave amplitude, holding covariates fixed. Positive coefficients indicate higher cognitive scores, and vice versa. For SZ, fitting the same equation to a 0/1 target gives a linear probability model. The amplitude coefficient represents the change in the modeled conditional probability of SZ per one-unit increase in amplitude. A positive coefficient indicates a higher SZ probability with higher amplitude; a negative coefficient indicates a lower probability.

#### 2) Phase-delay associations

To respect the circular scale of phase-delay, its sine and cosine were entered jointly^76^:

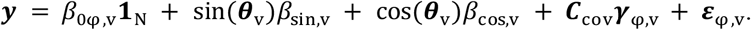

Here, **θ**_*v*_ ∈ ℝ^*N*×1^ contains one phase angle in radians at location v. Sine and cosine are applied on phase angle. The intercept *β*_0φ,v_ and coefficients *β*_sin,v_ and *β*_cos,v_ are scalars; ***γ***_φ,v_ ∈ ℝ^*Q*×1^ is the nuisance coefficient vector, the residual ***ε***_φ,v_ ∈ ℝ^*N*×1^ . Phase-delay association was tested by a joint F-test under the joint null hypothesis *H*_0_: *β*_sin,v_ = *β*_cos,v_ = 0. The joint F statistic compared the full model with a reduced model containing only the intercept and covariates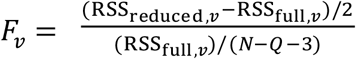 . RSS represents residual sum of squares, N, Q for sample number and covariate number. The numerator and denominator degrees of freedom were 2 and N-Q-3, respectively.

This tests whether phase is associated with cognitive score in HCP or the probability of SZ after covariate adjustment. Since the joint F statistic is nonnegative and has no directional sign. a significant joint test does not by itself establish that phase delays are shorter or longer with a phenotype. Directional analysis requires a separate phase-delay contrast between different phenotype groups.

To assess amplitude and phase-delay associations, P values were corrected across spatial locations using the Benjamini-Hochberg FDR procedure, separately for amplitude and phase analyses. Locations with FDR-adjusted P<0.05 were considered statistically significant.

## Acknowledgements

No competing interests.

## Data availability

The Human Connectome Project datasets are publicly available from the HCP-YA S1200 (https://www.humanconnectome.org/study/hcp-young-adult) hosted on the NIMH Data Archive (NDA) (https://nda.nih.gov/). The Functional Imaging Biomedical Informatics Research Network (FBIRN) Phase III dataset analyzed in this study is publicly available through the FBIRN resource hosted by the Neuroimaging Tools and Resources Collaboratory (NITRC) (https://www.nitrc.org/projects/fbirn/).

## Code availability

The pattern distribution of cortex and subcortex was visualized using the Nilearn and Pyvista Python packages. Code for simulation generation and related analyses could be made available in the supplementary files and shared on GitHub upon acceptance. The CPCA implementation used in this study was adapted from (https://github.com/facebookarchive/fbpca, as well as https://github.com/tsb46/complex_pca ). Software to obtain individual-specific network spatial patterns using spatial ICA and spatial-constrained ICA is available in the Group ICA of fMRI toolbox (GIFT) (https://trendscenter.org/software/). The PhaseICA pipeline is also available on the TReNDS GitHub site (https://github.com/trendscenter).

## Notes

### Competing Interest Statement

The authors have declared no competing interest.

### Summary of Updates

give me details of PhaseICA pipeline and application

